# Changes in both top-down and bottom-up effective connectivity drive visual hallucinations in Parkinson’s disease

**DOI:** 10.1101/2022.03.25.485621

**Authors:** George EC Thomas, Peter Zeidman, Tajwar Sultana, Angeliki Zarkali, Adeel Razi, Rimona S Weil

**Affiliations:** Dementia Research Centre, UCL Institute of Neurology, London, UK; Wellcome Centre for Human Neuroimaging, UCL, London, UK; Department of Computer and Information Systems Engineering, NED University of Engineering & Technology, Karachi; Department of Biomedical Engineering, NED University of Engineering & Technology, Karachi, Pakistan; Neurocomputation Laboratory, National Centre of Artificial Intelligence, Pakistan; Turner Institute for Brain and Mental Health, School of Psychological Sciences, Monash University, Clayton, Australia; CIFAR Azrieli Global Scholars Program, CIFAR, Toronto, Canada; Movement Disorders Consortium, UCL, London, UK

**Author notes:** **Corresponding author:** Mr George EC Thomas, Dementia Research Centre, University College London, 8-11 Queen Square, WC1N 3AR.

**Keywords:** Spectral dynamic causal modelling, resting-state functional MRI, Parkinson’s disease, visual hallucinations, effective connectivity

## Abstract

Visual hallucinations are common in Parkinson’s disease and are associated with poorer quality of life and higher risk of dementia. An important and influential model that is widely accepted as an explanation for the mechanism of visual hallucinations in Parkinson’s disease and other Lewy-body diseases is that these arise due to aberrant hierarchical processing, with impaired bottom-up integration of sensory information and overweighting of top-down perceptual priors within the visual system. This hypothesis has been driven by behavioural data and supported indirectly by observations derived from regional activation and correlational measures using neuroimaging. However, until now, there was no evidence from neuroimaging for differences in causal influences between brain regions measured in patients with Parkinson’s hallucinations. This is in part because previous resting-state studies focus on functional connectivity, which is inherently undirected in nature and cannot test hypotheses about directionality of connectivity. Spectral dynamic causal modelling is a Bayesian framework that allows the inference of effective connectivity – defined as the directed (causal) influence that one region exerts on another region – from resting-state functional MRI data. In the current study, we utilise spectral dynamic causal modelling to estimate effective connectivity within the resting-state visual network in our cohort of 15 Parkinson’s disease visual hallucinators, and 75 Parkinson’s disease non-hallucinators. We find that visual hallucinators display decreased bottom-up effective connectivity from the lateral geniculate nucleus to primary visual cortex and increased top-down effective connectivity from left prefrontal cortex to primary visual cortex and medial thalamus, as compared to non-hallucinators. Importantly, we find that the pattern of effective connectivity is predictive of the presence of visual hallucinations and associated with their severity within the hallucinating group. This is the first study to provide evidence, using resting state effective connectivity, to support a model of aberrant hierarchical predictive processing as the mechanism for visual hallucinations in Parkinson’s disease.

## INTRODUCTION

Visual hallucinations (VH) are common in Parkinson’s disease, affecting up to 70% of patients^1^. Parkinson’s disease patients who suffer from VH have a poorer quality of life^2^, an increased rate of mortality^3^, and are more likely to require nursing home care^4^ and to develop subsequent dementia^5^. Until recently, models for how VH are generated within the brain remained somewhat limited and disparate, despite the high prevalence of VH and the clear burden they have on patients and their carers^6^.

However, over the last few years, lines of evidence have converged on a predictive-coding framework that suggests the occurrence of VH relates to altered integration of sensory input (bottom-up information) and prior knowledge (top-down information, or a ‘world model’) within the visual system^7,8^. Patients with Parkinson’s disease frequently have deficits in visual processing^6,9–12^, with potentially greater deficits in higher-order visual processing in Parkinson’s patients who hallucinate compared with those that do not^13^. Behavioural evidence has pointed specifically to impaired bottom-up accumulation of sensory evidence in visual hallucinators as compared to non-hallucinators^14^, and patients with Lewy-body related VH have been shown to rely more strongly on top-down perceptual priors^15^. Notably, dorsolateral regions of prefrontal cortex are thought to play a key role in controlling top-down causal prediction error signalling^16,17^.

Several key brain regions have been implicated in Parkinson’s hallucinations. The thalamus is expected to be important in this framework as the synchronised functioning of these systems during normal visual perception relies strongly on thalamocortical connections^18,19^. Previous work has shown that Parkinson’s disease hallucinators have reduced fibre cross section in white matter tracts connected to the medial thalamus early in disease course, with this spreading to almost the whole thalamus in later disease^20^. Additionally, lesion network mapping has revealed that a network centred on the lateral geniculate nucleus is consistently affected in studies of Parkinson’s hallucinations^21^ and in patients with brain lesions causing visual hallucinations^22^.

Evidence also points towards the hippocampus as being important for application of perceptual priors due to its role in the encoding and retrieval of event-related memories^23^ as well as the integration of spatial and non-spatial contextual information^24^. In Parkinson’s disease, VH have been linked to a higher burden of Lewy-related pathology in medial temporal lobes^13,25^. Alterations in functional connectivity to the hippocampus is seen in Parkinson’s hallucinators, with increased functional connectivity to default mode and frontal regions, and decreased functional connectivity with the occipital cortex ^26^.

Primary visual cortex and frontal regions have been implicated in numerous functional MRI studies of Parkinson’s hallucinators^27^. During exposure to simple visual stimuli, decreased activation in the primary visual and occipital cortex^28–30^, has frequently been observed alongside increased activation in frontal^28,30^ and visual association regions^29^ in visual hallucinators. Exposure to complex visual stimuli in Parkinson’s disease with VH is associated with decreased frontal cortical activation^31,32^ and with altered functional connectivity in and between the attentional and default mode networks^33^.

Altered functional connectivity has also been reported in resting-state analyses comparing Parkinson’s disease visual hallucinators with non-hallucinators^27^. Studies looking at the default mode network have found mostly increased functional connectivity in hallucinators compared to non-hallucinators, including in frontal-parietal regions^34,35^. Increased connectivity within and between the default mode and attentional networks has also been found to correlate with performance on a complex visual task^36^. Hallucination severity is associated with strengthened stability of the default mode network^37^, and greater mind wandering in visual hallucinators linked with increased functional connectivity between primary visual cortex and the dorsal default mode network^38^.

Outside of the default mode network, increased occipital functional connectivity has been described with frontal and cortico-striatal regions^39^ and decreased functional connectivity has been reported between the posterior cingulate cortex and parietal, temporal and occipital regions^40^ in Parkinson’s disease with VH.

Whilst these studies confirm altered connectivity and activation within visual network regions relating to VH in Parkinson’s disease, these studies only examine differences in functional connectivity, which is a measure of the statistical dependencies or correlations between neuroimaging timeseries. The correlational, undirected nature of functional connectivity precludes any assessment of the *causal* influences of distinct brain regions and cannot provide a mechanistic explanation of neural mechanisms of Parkinson’s hallucinations based on hierarchical predictive processing which requires information on directionality to address the question of relative importance of bottom-up and top-down signalling.

Here, we employ dynamic causal modelling - a Bayesian framework that allows us to infer effective connectivity, which is designated as the directed (causal) influences among brain regions ^41^. A dynamic causal model (DCM) is defined by a forward model that generates neuroimaging timeseries based on the underlying causes, controlled by the model parameters. These parameters represent quantities such as connection strengths between regions, and fall into three categories: neuronal parameters, haemodynamic parameters and parameters due to noise or measurement error^42^. Once a DCM is specified, data can be simulated under different variations of the model (for example with different connection strengths or architectures) to determine which model best characterises the observed data. Hypotheses within or between subjects can then be tested by comparing the evidence for different models.

When modelling resting state functional MRI data, the DCM methodology can become computationally intensive, due to the need to estimate random fluctuations in neuronal states^43^. Spectral DCM^41^ has the advantage over conventional DCM when applied to resting state data, in that rather than modelling time-varying fluctuations in neuronal states, it models their second-order statistics or cross-spectra. This essentially models the dynamics of different brain regions in the frequency domain rather than the time domain. In doing so, it eliminates the need to estimate random neuronal fluctuations and is therefore more computationally efficient^44^. Additionally, it provides a reliable estimation of between region influences as well as enabling the accurate detection of group differences in effective connectivity.

In the current study, we use spectral dynamic causal modelling to estimate effective connectivity within a set of pre-defined visual brain regions in Parkinson’s disease patients with and without visual hallucinations. We investigate the importance of top-down and bottom-up connectivity in explaining the differences between these groups, as well as the relative contributions of different regions and hemispheric connections to provide information about the architecture of this connectivity. We hypothesised that the presence of VH would be associated with both reduced bottom-up and increased top-down connectivity within this network, and that the severity of hallucinations would be associated with the pattern of changes in connectivity.

## RESULTS

### Participants

Of the 90 PD patients in the current study, 15 were identified as having VH (PD-VH), while 75 had no VH (PD-no-VH). The PD-no-VH group had a higher female to male ratio than the PD-VH group (T=3.90, p=0.045), but there were no significant differences in the other demographics or clinical metrics, including no difference in measures of cognition or Levodopa equivalent dose (see Table 1).

**Table 1.** Participant demographics. Means (SDs) reported. MoCA = Montreal Cognitive Assessment. UPDRS = Unified Parkinson’s Disease Rating Scale. HADS = Hospital Anxiety and Depression Scale. Binocular LogMAR: lower score is better visual acuity. RBDSQ = REM (Rapid Eye Movement) Sleep Behaviour Disorder Screening Questionnaire. *P<0.05; ns = not significant

| Measure | PD-no-VH<br>(N=75) | PD-VH<br>(N=15) | Statistic | *P* |  |
| --- | --- | --- | --- | --- | --- |
| Sex (M:F) | 45:31 | 4:11 | $T = 3.90$ | 0.045 | * |
| Age (years) | 64.12 (7.78) | 65.33 (8.75) | $OR = -0.10$ | 0.92 | ns |
| Years of education | 16.83 (2.73) | 17.77 (3.59) | $U = 3311$ | 0.27 | ns |
| MoCA (out of 30) | 28.15 (2.08) | 27.60 (1.76) | $U = 3546$ | 0.14 | ns |
| UPDRS-III | 21.89 (11.34) | 25.40 (14.67) | $U = 3331$ | 0.38 | ns |
| Binocular LogMAR visual acuity | -0.09 (0.12) | -0.04 (0.34) | $U = 3325$ | 0.34 | ns |
| HADS depression score | 3.97 (2.95) | 4.73 (3.43) | $U = 3334$ | 0.39 | ns |
| HADS anxiety score | 5.77 (3.80) | 7.00 (4.38) | $U = 3314$ | 0.29 | ns |
| RBDSQ score | 3.96 (2.41) | 5.13 (2.53) | $U = 3255$ | 0.08 | ns |
| Smell test (Sniffin' Sticks) | 7.92 (3.16) | 6.40 (3.29) | $T = 1.69$ | 0.09 | ns |
| Disease duration (years) | 3.83 (2.38) | 4.67 (2.35) | $U = 3286$ | 0.17 | ns |
| Levodopa equivalent dose (mg) | 431.89<br>(259.88) | 421.67<br>(196.57) | $U = 3389$ | 0.80 | ns |

### Accuracy of DCM model estimation

The variance explained by DCM model estimation when fitted to the observed spectral data was 89.6±3.3% (mean ± standard deviation), with a range of 81.22% to 96.96%. This confirms the good fits of estimated DCM to the empirical cross-spectra. An example participant’s estimation across spectral densities and estimated connectivity parameters can be seen in Figure 1C-D. Cross-spectral density plots for all 15 PD-VH and a summary plot across all subjects can be seen in supplementary figure 1.

**Figure 1.**
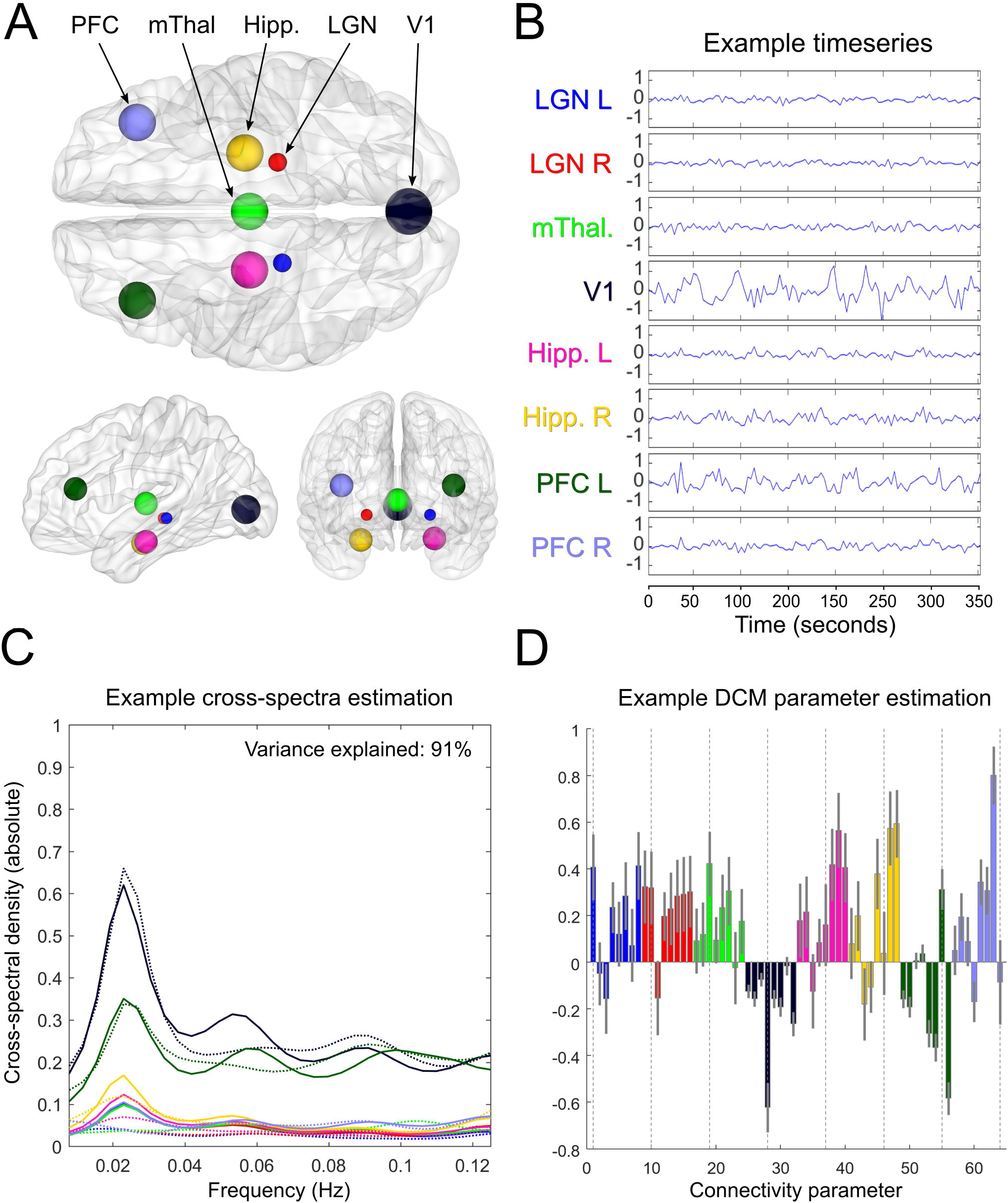
Within-subject dynamic causal modelling analysis. **(A)** Location of the eight nodes comprising the visual network used for all analyses (Dorsal, left-sagittal, and rostral views). **(B)** Timeseries extracted as the principal eigenvariate from each of the eight visual network nodes (example from a Parkinson’s hallucinator, data from same patient shown for C and D) **(C)** Cross-spectral density plot, showing the frequency bands in which each of the eight regions were active. Real data are indicated by solid lines (and are second order statistics derived from the timeseries), while data estimated by the generative model are indicated by dashed lines. The variance explained by the model for this example subject is indicated in the top right. **(D)** Estimated connectivity parameters (connection strengths) for the eight regions, including 56 extrinsic (between-region) parameters, which are in hertz, and eight intrinsic (within-region) parameters indicated by dotted lines, which have a unitless log scale and modulate inherently negative self-connections. Error bars are 90% credible intervals derived from the posterior variance of each parameter. LGN = lateral geniculate nucleus; mThal. = medial thalamus; V1 = primary visual cortex; Hipp. = hippocampus; PFC = prefrontal cortex; DCM = dynamic causal model. Colour coding for regions is consistent across panels.

### Commonalities of visual network effective connectivity across participants

The visual network architecture common to all participants with Parkinson’s disease was defined by the commonalities in connection strengths, or group mean, identified in our hypothesis-based analysis (hypothesis-based analysis detailed in Figure 2). Across all reduced GLMs, the highest associated posterior probability was 53% (Figure 3A). This GLM had its commonalities quantified by the effective connectivity parameters of model 172 (Figure 3C, left panel), which was the fully connected model (all connections switched on). Summing across all GLMs which had their commonalities parameters deployed according to this model, the posterior probability reached 99.8% (Figure 3B, upper panel). This strong evidence indicates that the commonalities across subjects were best explained by the fully connected model. Thus, no connections could be excluded without reducing the quality of the model. Accordingly, family-based analyses revealed that both top-down and bottom-up connections, (family 3 in factor 1, Figure 3D), both inter- and intra-hemispheric connections (family 3 in factor 2, Figure 3E), and connections to and from all regions (family 23 in factor 3, Figure 3F) were important in explaining the commonalities across subjects.

**Figure 2.**
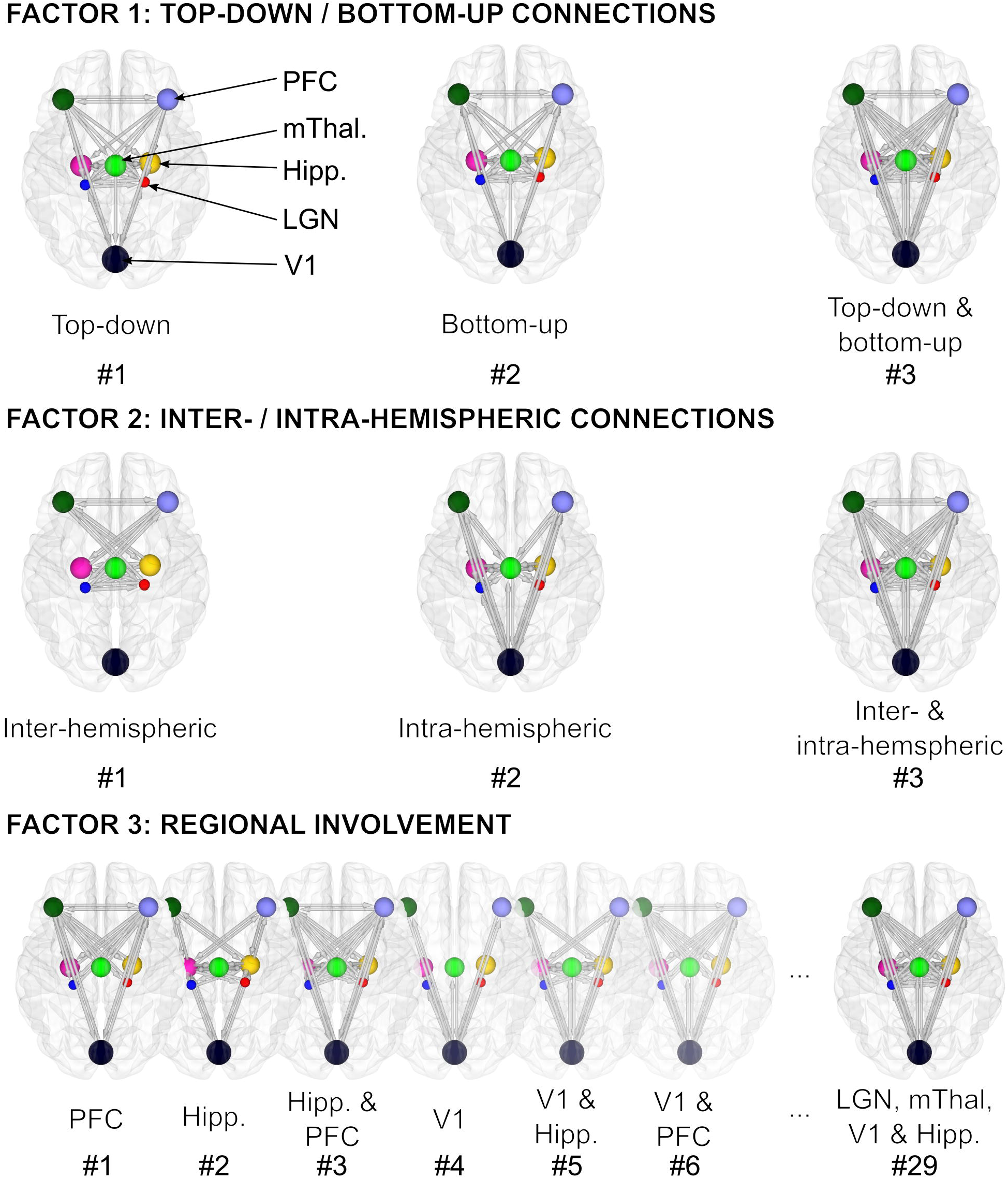
Construction of factorial model space to test hypotheses about visual network hierarchical processing and architecture in Parkinson’s disease with visual hallucinations. Possible explanations for the data in the form of reduced models (i.e., some connections ‘switched off’ / restricted to their prior expectation of zero) were constructed based on their correspondence to three experimental factors for both the commonalities and differences between subjects. **Factor 1** examines top-down vs bottom-up connectivity and has 3 families within it: 1) – **Top-down:** bottom-up connections switched off, 2) – **Bottom-up:** top-down connections switched off and 3) – **Top-down and bottom-up:** no connections switched off. **Factor 2** examines inter-vs intra-hemispheric connectivity and has 3 families within it: 1) - **Inter-hemispheric:** intra-hemispheric connections switched off, 2) **Intra-hemispheric** inter-hemispheric connections switched off and 3) **Inter-hemispheric** and **Intra-hemispheric:** no connections switched off. **Factor 3** examines regional involvement and has 29 families within it, which were derived from all possible subsets of five bilateral regions (left and right LGN, medial thalamus, V1, left and right hippocampus, left and right PFC). For example, #1 all extrinsic connections switched off except those to and from the left and right PFC, #29 all extrinsic connections switched on except those specific to left and right PFC. All possible familial combinations across factors gave 3×3×29 = 261 possible models. After removal of duplicate models and addition of a null model with no connections switched on (including intrinsic ones), the final factorial model space contained 179 models each for the commonalities and differences between subjects. Therefore, 179^2^ = 32041 possible hypotheses (PEB models) for the data were analysed.

**Figure 3.**
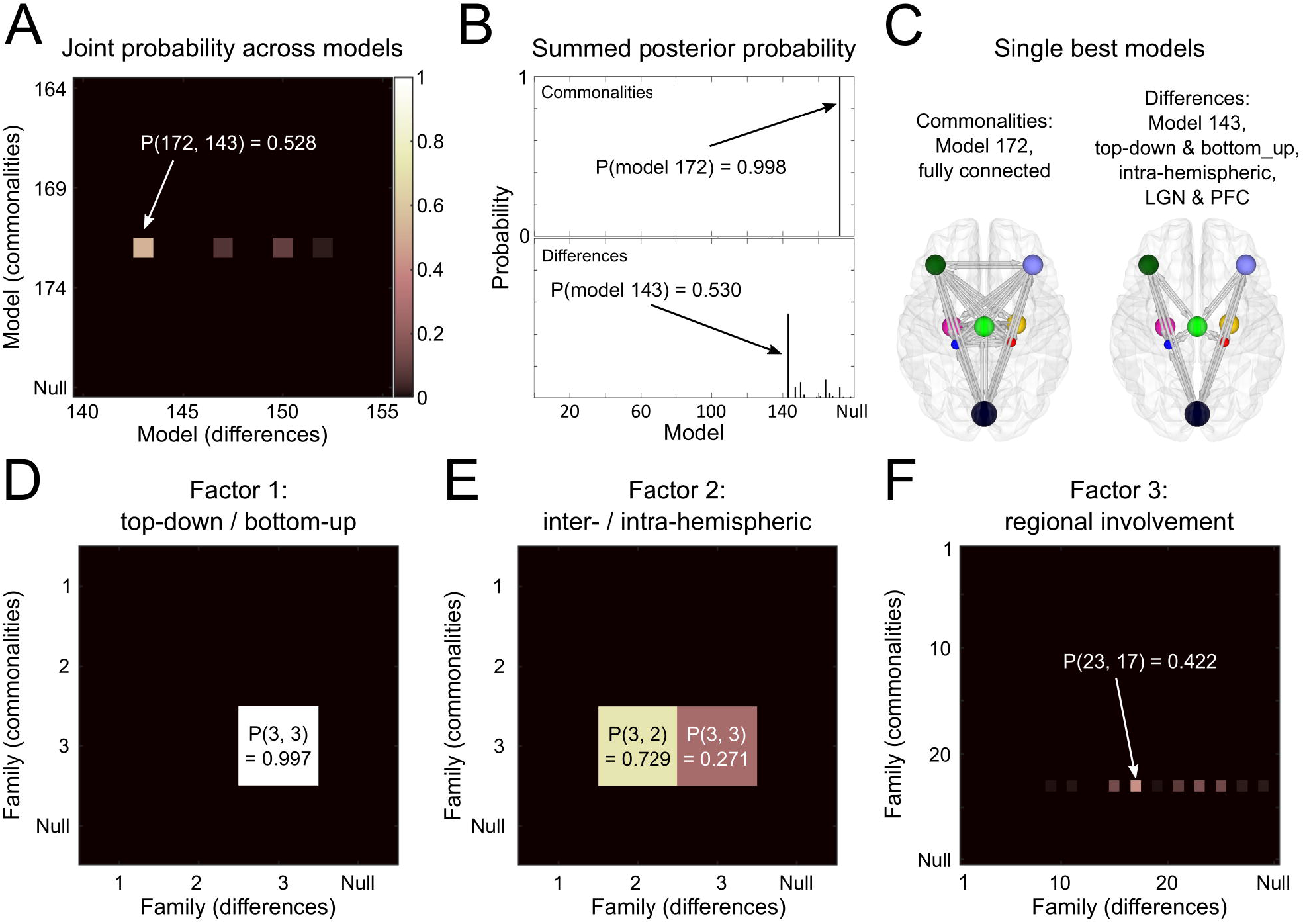
Bayesian model comparison across factorial model space gives possible explanations for visual network hierarchical processing and architecture. **(A) Joint posterior probability across all models**. The axes list all 179 candidate models including the null model in terms of the commonalities across subjects and differences due to presence of visual hallucinations (i.e., the hypothesis in row *i* and column *j* had parameters relating to commonalities set according to model *i*, and parameters relating to presence of hallucinations set according to model *j*). The best model was number 172 (fully connected) for the commonalities and 143 (top-down and bottom-up with intra-hemispheric LGN and PGC) for differences, with 53% posterior probability. **(B) Summed posterior probability**. The same result shown in panel A, summed over the columns to give the posterior probability for the commonalities across subjects, and summed over the rows and to give the posterior probability for the differences due to hallucinations. **(C)** Spatial projections for the two single-best models described in panels A and B. **(D-F) Family analyses for each factor: D: top-down/bottom-up; E inter-/intra-hemispheric; F regional involvement**. Separate family analyses with the same 32041 hypothesis (reduced PEB models) from panel A grouped into different families. In each case, the element in row *m* and column *n* represents the pooled probability across models in which the commonalities parameters were set according to family *m* and the parameters relating to presence of hallucinations set according to family *n*. Factors and families are as described in Figure 2, and the probability associated with the null model is also plotted in each case.

Bayesian parameter averaging of commonalities between subjects revealed that this fully connected model was characterised by strongly positive interhemispheric effective connectivity, and strongly positive intrinsic (self) connectivity in all regions except the PFC and medial thalamus. Note that effective connectivity with a positive sign represents excitatory influences, and negative sign represents inhibitory influences except for (log-scaled) self-connections which are always inhibitory by definition. Hence positive self-connections represent more inhibition and negative self-connections represent disinhibition. Other key features included generally negative afferent PFC connectivity, and positive connectivity to V1 from LGN and PFC (Figure 4A). This architecture was confirmed by our data-driven automatic search over parameters, which identified a highly connected network with very similar connection strengths (supplementary figure 2).

**Figure 4.**
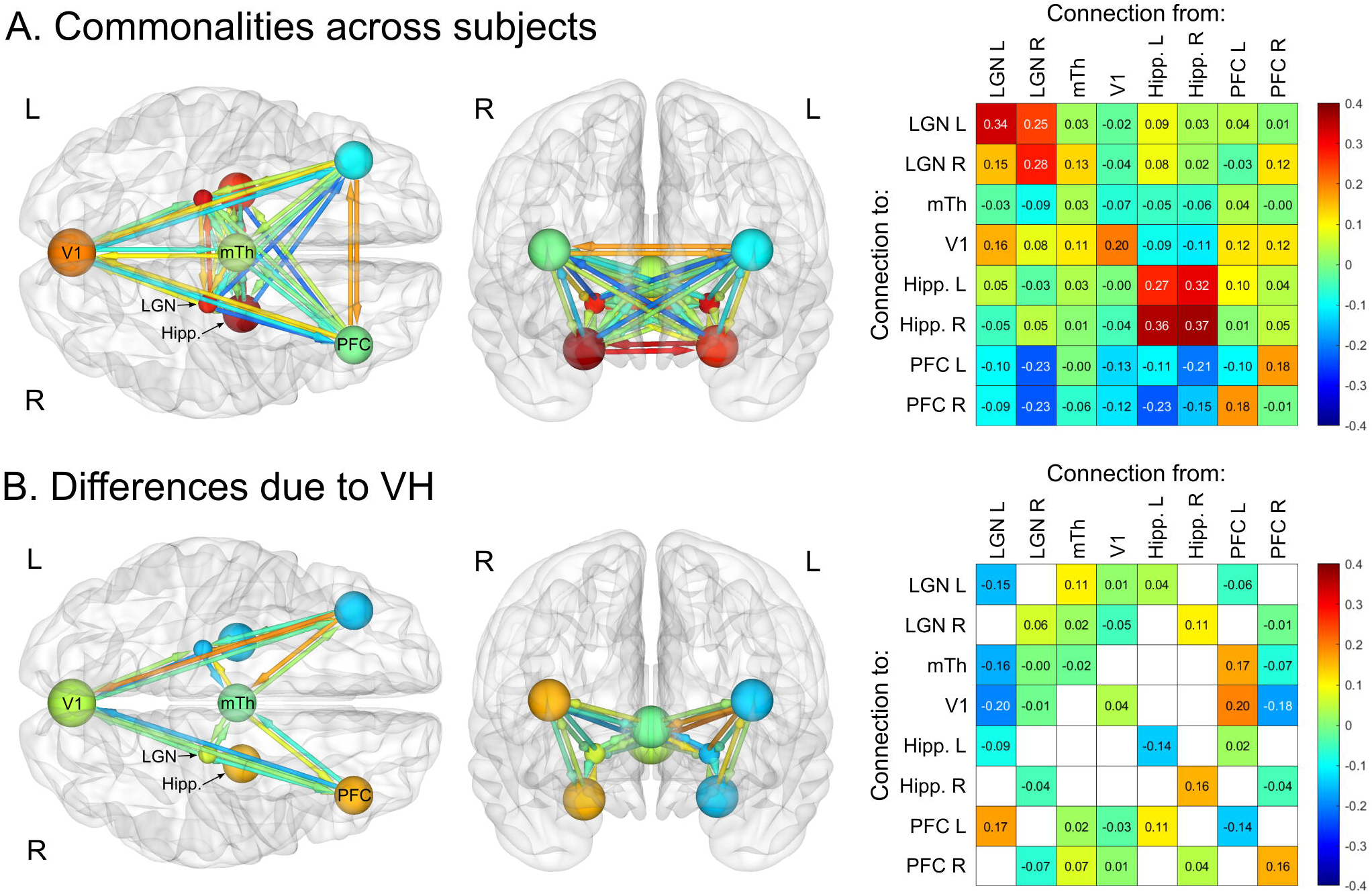
Bayesian model averaging across factorial model space gives magnitude of connection strengths across subjects, and differences associated with visual hallucinations. The Bayesian model average of parameter values across all 179 models for the commonalities between subjects and the differences between subjects due to the presence of visual hallucinations. All parameters are thresholded at posterior probability >95% of being present versus absent. **(A) Commonalities across all patients** (with and without hallucinations equivalent to the mean across subjects). Here, orange arrows / off-diagonal positive numbers reflect excitatory connectivity and blue arrows / off-diagonal negative numbers reflect inhibitory connectivity. Diagonal connectivity / self-connectivity is inhibitory by definition and is log-scale, hence leading diagonal positive numbers / orange spheres reflect more self-inhibition and leading diagonal negative numbers / blue spheres reflect disinhibition. **(B) Differences between hallucinators and non-hallucinators**. Here, orange arrows / off-diagonal positive numbers reflect increased connectivity in PD-VH versus PD-no-VH, whereas blue arrows / off-diagonal negative numbers reflect decreased connectivity. Leading diagonal positive numbers / orange spheres reflect increased self-inhibition in PD-VH versus PD-no-VH and leading diagonal negative numbers / blue spheres reflect increased disinhibition. LGN = Lateral geniculate nucleus; mTh = medial thalamus; Hipp. = hippocampus; PFC = prefrontal cortex.

### Effect of visual hallucinations on visual network architecture

The differences in visual network architecture in PD-VH compared to PD-no-VH were best explained by GLMs with parameters deployed according to model 143 (Figure 3A), with an associated summed posterior probability of 53% (Figure 3B bottom). This model was defined by the presence of both top-down and bottom-up intra-hemispheric connections to and from LGN and PFC (Figure 3C, right panel). Accordingly, family-based analysis revealed that both top-down and bottom-up connections, (family 3 in factor 1, Figure 3D), intra-hemispheric connections (family 2 in factor 2, Figure 3E), and connections to and from LGN and PFC (family 17 in factor 3, Figure 3F) were the best explanation for differences between subjects due to presence of VH.

While model 143 had the strongest posterior probability across all 179 models, its overall probability was only 53%. There was therefore not a single best explanation for the differences in connectivity due to VH (i.e., a model with 95% probability or more^45,46^). This *dilution of evidence* effect is expected given the large number of models. Therefore, to summarise the estimated parameters across the 32041 models, while taking into account that different models had different levels of evidence, we performed Bayesian model averaging. We then thresholded the averaged parameters at >95% posterior probability, corresponding to strong evidence^45,46^, for effects being present versus absent.

Bayesian model averaging across all models in the factorial model space revealed several key differences in effective connectivity in PD-VH compared to PD-no-VH. These included reduced effective connectivity from left LGN to V1 (0.20Hz) and medial thalamus (0.16Hz), increased effective connectivity from left LGN to left PFC (0.17Hz), increased effective connectivity from left PFC to medial thalamus (0.17Hz) and V1 (0.20Hz) and decreased effective connectivity from right PFC to V1 (0.18Hz). Additionally, in PD-VH, we observed increased self-inhibition in the right PFC and right hippocampus, and disinhibition of the left PFC, left hippocampus and left LGN. The full pattern of connectivity differences can be seen in Figure 4B. The differences between subjects due to the presence of VH identified after an automatic search over parameters were in concordance with those described above (supplementary figure 2).

This indicates that both increased top-down and reduced bottom-up effective connectivity is important in explaining the differences between PD-VH and PD-no-VH groups, particularly relating to the LGN and PFC, with a lateralised effect observed from the PFC.

### Predicting group membership and hallucination severity

We used leave-one-out cross validation to test whether a participant’s group membership (i.e., hallucination status) could be predicted from their individual connection strengths for the top five largest group differences, as previously descried in the Methods section. We found that 71 out of 90 subjects had their true group membership value within the estimated 90% credible interval, with 14 out of 15 PD-VH and 57 out of 75 PD-no-VH falling in this range. Predicted group membership correlated significantly with actual group membership (r = 0.24, *P* = 0.012; Figure 5A). Additionally, this correlation remained significant when predicting group membership using the top 10 connections (r = 0.30, *P* = 0.002) or the single largest connection (r = 0.22, *P* = 0.012).

**Figure 5.**
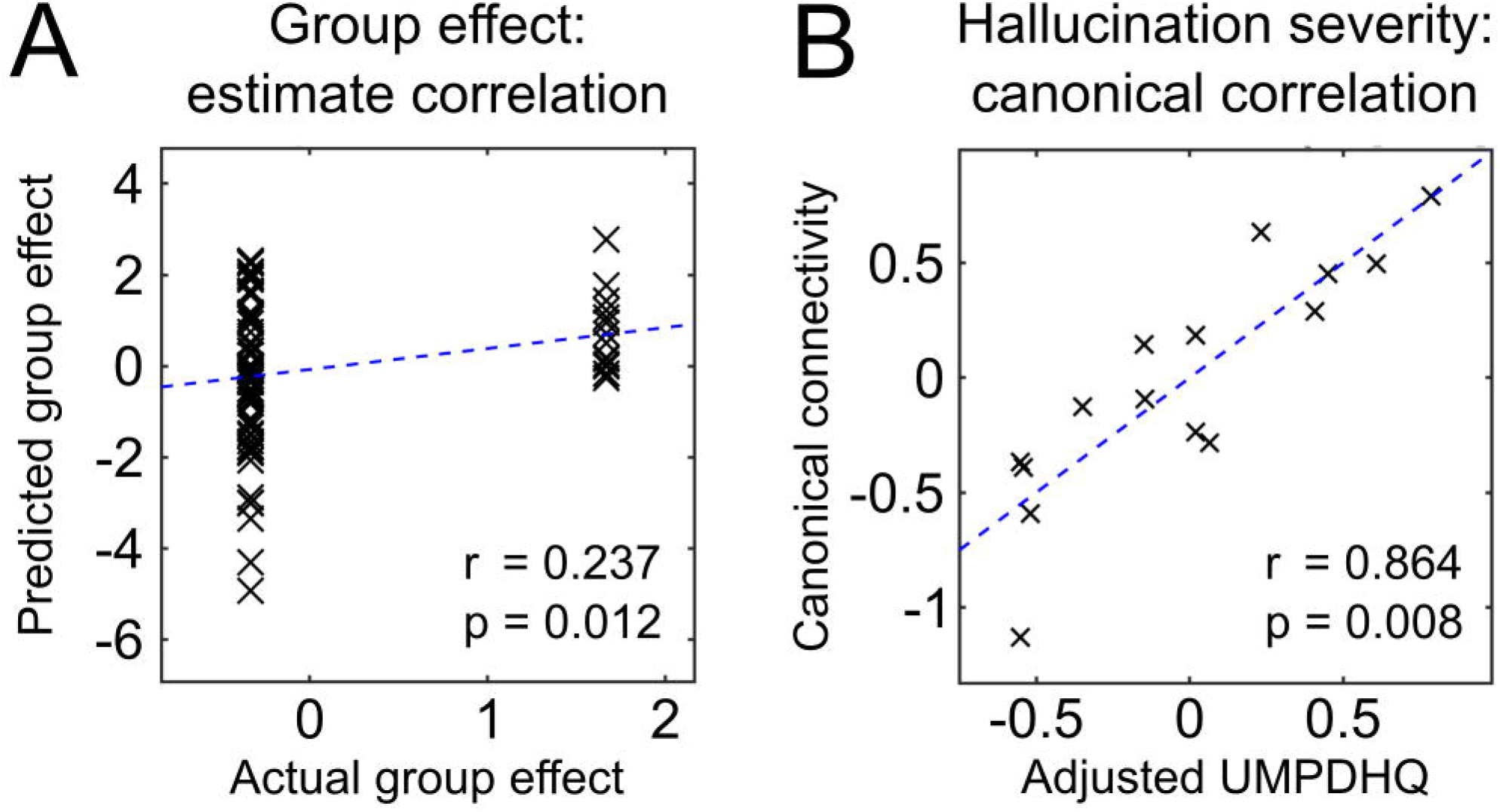
Visual network effective connectivity is associated with the presence and severity of visual hallucinations in Parkinson’s disease. **(A) Results from leave-one-out cross validation across all 90 subjects**. The five largest parameters in terms of overall group difference were used to predict the left-out subject’s group membership. Correlation between predicted group effect and actual group effect, r is the Pearson’s correlation coefficient. **(B) Relationship between differences in effective connectivity and hallucination severity**. Results of canonical variate analysis for the 15 PD-VH. For each participant, the primary canonical variate derived from their individual values for the five largest parameters in terms of overall group difference between PD-VH and PD-no-VH is plotted against the primary canonical variate derived from their UMPDHQ score adjusted for age and sex, r is the Pearson’s correlation coefficient.

We next investigated whether hallucination severity within the PD-VH group was associated with the first canonical variate derived from five parameters described above. We found that this correlated significantly with the canonical variate derived from adjusted UMPDHQ score (r = 0.864, *P* = 0.008; Figure 5B), indicating that the pattern of connectivity within the visual network in the PD-VH group was predictive of hallucination severity. This association remained significant when using the top 10 connection strengths (r = 0.854, *P*=0.018) and approached significance when using the single largest connection (r = 0.643, *P*=0.053).

## DISCUSSION

In the current study, we utilised spectral DCM for resting state functional MRI data to investigate visual network effective connectivity in Parkinson’s disease patients with and without VH. We examined how effective connectivity differed between patients with and without VH, finding that a model comprising both reduced bottom-up and increased top-down effective connectivity best explained differences between patient groups, with differences in effective connectivity relating to severity of hallucinations. To our knowledge, this is the first study demonstrating changes in causal influences between brain regions in Parkinson’s disease visual hallucinators.

The ability of DCM to non-invasively infer the directionality of influences between brain regions in living patients^47^ ideally places it to investigate hypotheses relating to the hierarchical processing account of VH, where reduced bottom-up sensory processing is proposed to occur alongside overweighting of top-down perceptual priors^14,15^. Previous studies, while making valuable insights^27,48^, have tended to look at functional connectivity changes, which are inherently undirected in nature^47^, and so cannot directly test this hypothesis. We designed a factorial model space to specifically probe the relative contribution of top-down and bottom-up connections within the visual network in PD-VH versus PD-no-VH, and additionally tested inter-versus intra-hemispheric connectivity alongside regional involvement to estimate the architecture of any changes. Finally, we assessed whether the pattern of visual network changes observed was associated with a clinically-relevant measure: the severity of hallucinations.

Our Bayesian model comparison across this model space revealed that the differences between subjects due to the presence of VH were explained best by a combination of both top-down and bottom-up effective connections. In particular, these were intra-hemispheric connections to and from the LGN and PFC. Bayesian model averaging across the model space revealed both the valence and magnitude of such changes. Among the largest were decreased effective connectivity in PD-VH from the left LGN to medial thalamus and V1, as well as increased effective connectivity from left PFC to medial thalamus and V1. Thus, some of the largest differences we found between PD-VH and PD-no-VH reflected both increased top-down and reduced bottom-up connections within the visual network, with regions at either end of the visual hierarchy (LGN and PFC) emerging as particularly important.

The single largest difference we found between PD-VH and PD-no-VH was decreased effective connectivity from left LGN to V1 in PD-VH. This is consistent with previous structural network mapping work, which found that coordinates of atrophy from studies of VH in Parkinson’s disease were connected to a network centred on the LGN^21^. Similarly lesions in patients with visual hallucinations are connected to an LGN-centred functional network^22^. Task-based functional MRI work in PD-VH has also shown decreased activity in primary visual and occipital cortex fairly consistently in PD-VH compared to PD-no-VH^28–30^. Additionally, magnetic resonance spectroscopy has revealed reduced occipital GABA^49^ and FDG-PET has revealed occipital glucose hypometabolism associated with VH in Parkinson’s disease^50,51^. Reduced activity of early visual cortex would be consistent with reduced bottom-up input from LGN, although we did not find altered efferent connectivity from V1 in PD-VH. Interestingly, a recent study of effective connectivity in Parkinson’s disease without VH found that modulation of LGN by the superior colliculus in response to luminance contrast changes was inhibited in Parkinson’s disease compared to controls^52^. The way in which such stimuli modulate LGN and the superior colliculus may also change as a result of treatment^53^.

We observed changes in effective connectivity to the medial thalamus in PD-VH, with decreased connectivity from the left LGN, and increased connectivity from left PFC. This is congruent with our recent work finding that thalamic tracts connected to the mediodorsal thalamic nucleus had reduced fibre cross section in PD-VH^20^, although these findings did not indicate the direction of information flow. Previous diffusion tensor imaging work revealed increased mean diffusivity in bilateral thalamic sub regions projecting to prefrontal and parieto-occipital cortices in Lewy body dementia patients with VH^54^. Additionally, a recent post-mortem study found that atrophy in the mediodorsal nucleus of the thalamus was significantly greater in Lewy body dementia patients with VH^55^.

The thalamus is expected to play an important role in accounts of dysfunctional hierarchical predictive processing, due to its role in synchronising simultaneous streams of information (e.g. top-down and bottom-up) in normal visual processing^18,19^. This is corroborated by the changes in thalamic effective connectivity we found in PD-VH. Interactions between thalamus and prefrontal cortex are important in dealing with perceptual uncertainty during decision making^56^. Two separate pathways may exist for this, with low-signal-related uncertainty (as would be expected with reduced bottom-up sensory input) resolved by dopaminergic projections from medial thalamus that increase prefrontal output^57^. Interestingly, changes in thalamic resting state functional connectivity^58^, as well as changes in thalamocortical effective connectivity^59^ may also underlie hallucinations during exposure to LSD, consistent with a more general role for changes in thalamic connectivity in hallucinations in other contexts^60^.

We also found increased top-down effective connectivity in PD-VH from left PFC to medial thalamus, and V1. This might help to explain previous task-based functional studies reporting increased activity in PD-VH prefrontal cortices in response to simple stimuli^28,30^, although other work has shown that prefrontal cortex activity may be decreased in PD-VH vs PD-no-VH in response to more complex stimuli^31,32^. Similarly, previous resting state-work has described increased functional connectivity between superior, middle and inferior frontal gyri in PD-VH and occipital cortex^39^. An increased burden of Lewy-related pathology has also been reported in frontal cortex in PD-VH^13^, as have grey matter reductions in the bilateral dorsolateral PFC^61^.

The lateralised effect of PFC efferent connectivity we found is intriguing, in that we found increased top-down connectivity from left PFC to V1 and medial thalamus, but decreased top-down connectivity from right PFC to V1. Lateralised differences in prefrontal activity have also been reported in previous functional neuroimaging studies of PD-VH. Both increased^50^ and decreased^51^ glucose metabolism have been observed in left PFC. Decreased right PFC BOLD activation has also been reported in PD-VH in response to faces^31^, and just prior to presentation of complex stimuli^32^. Increased right PFC BOLD activation has meanwhile been reported in response to more simple stimuli^30^. A single-case study has also reported simultaneous activation of right medial frontal gyrus and deactivation of left middle frontal gyrus during active VH^62^.

Asymmetry in lateral PFC influences have been shown in prediction error signalling in health, however right (rather than left) lateral PFC was implicated^16,17^. Our observation of increased effective connectivity from left PFC and decreased from right PFC is thus inconsistent with these findings. It is possible that Parkinson’s disease disrupts the usual laterality of prediction error signals, though this needs to be tested in larger numbers of patients.

Although we found evidence to support the increased top-down and reduced bottom-up connectivity account of VH in Parkinson’s disease, differences in effective connectivity between other regions are also implicated. For example, we also observed increased bottom-up connectivity from the left LGN and decreased top-down connectivity from right PFC in PD-VH, and this is further complicated by changes in regional self-connections. We found disinhibition of the left LGN (reduced intrinsic self-inhibition) which would be expected to amplify both the reduced bottom-up connectivity to medial thalamus and V1, and the increased bottom-up connectivity to left PFC. We further found disinhibition of the left PFC, which would amplify its increased top-down connections to medial thalamus and V1, and we found increased self-inhibition of right PFC, which would dampen its reduced top-down connection to V1. Of note, the combined differences in connectivity related to clinical measures of hallucination severity (both top 5 and top 10 connections) so a more complex shift in causal influences may be relevant. It is therefore important to consider the full set of changes in effective connectivity as well as both extrinsic and intrinsic connections when interpreting our results.

### Limitations

One of the main limitations with the current paper is the relatively small number of PD-VH included. While similar sample sizes have been used in previous functional analyses of VH in Parkinson’s disease^34,63^, it will be important to examine effective connectivity in larger cohorts.

Patients in the current study continued their normal medication, including levodopa, during both clinical assessments and neuroimaging. This was done to avoid potential effects of distress and anxiety caused by omitting levodopa doses, which would affect cognitive function. Although we are not able to directly assess the effect of neurotransmitters on the current results, we note there was no significant difference in levodopa equivalent dose between hallucinators and non-hallucinators.

Additionally, while predicted group membership correlated significantly with actual group membership in our leave-one-out analysis, we were not able to predict group membership on a per-subject basis with high posterior probability. This could be explained by heterogeneity within the PD-no-VH group, whereby those who will go on to develop VH show a similar pattern to those who already have VH. However, longitudinal data would be needed to test this. That said, we did find that the pattern of connectivity was associated with hallucination severity within the PD-VH group.

## Conclusions

Our spectral DCM analysis showed that VH in Parkinson’s disease is associated with both reduced bottom-up and increased top-down effective connectivity within the visual network. This related particularly to intra-hemispheric connectivity to and from the LGN and PFC. The pattern of effective connectivity within the PD-VH group was significantly associated with the severity of their hallucinations. This study provides further evidence for the aberrant hierarchical predictive processing account of hallucinations in Parkinson’s disease, and models resting state effective connectivity for the first time in this context.

## Abbreviations

CVA: canonical variates analysis
DCM: dynamic causal modelling
LGN: lateral geniculate nucleus
PEB: parametric empirical Bayes
D-no-VH: Parkinson’s disease without visual hallucinations
PD-VH: Parkinson’s disease with visual hallucinations
PFC: prefrontal cortex
V1: primary visual cortex
VH: visual hallucinations

## FUNDING

GECT is supported by a PhD studentship from the Medical Research Council (Ref: MR/N013867/1). PZ is supported by core funding awarded by Wellcome, to the Wellcome Centre for Human Neuroimaging (Ref: 203147/Z/16/Z). AZ is supported by an Alzheimer’s Research UK Clinical Research Fellowship (Ref: 2018B-001). AR is supported by the Australian Research Council (Refs: DE170100128 and DP200100757) and Australian National Health and Medical Research Council Investigator Grant (Ref: 1194910). AR is also a CIFAR Azrieli Global Scholar in the Brain, Mind & Consciousness Program. RSW is supported by a Wellcome Clinical Research Career Development Fellowship (Ref: 205167/Z/16/Z). Recruitment to the study was also supported by Parkinson’s UK, the Cure Parkinson’s Trust. The study was further supported by UCLH Biomedical Research Centre Grant (Ref: BRC302/NS/RW/101410) and by grants from the National Institute for Health Research.

## COMPETING INTERESTS

RSW has received honoraria from GE healthcare and Britannia.

