## Supplementary figures and images for "Changes in both top-down and bottom-up effective connectivity drive visual hallucinations in Parkinson’s disease"

### Supplementary figure 1

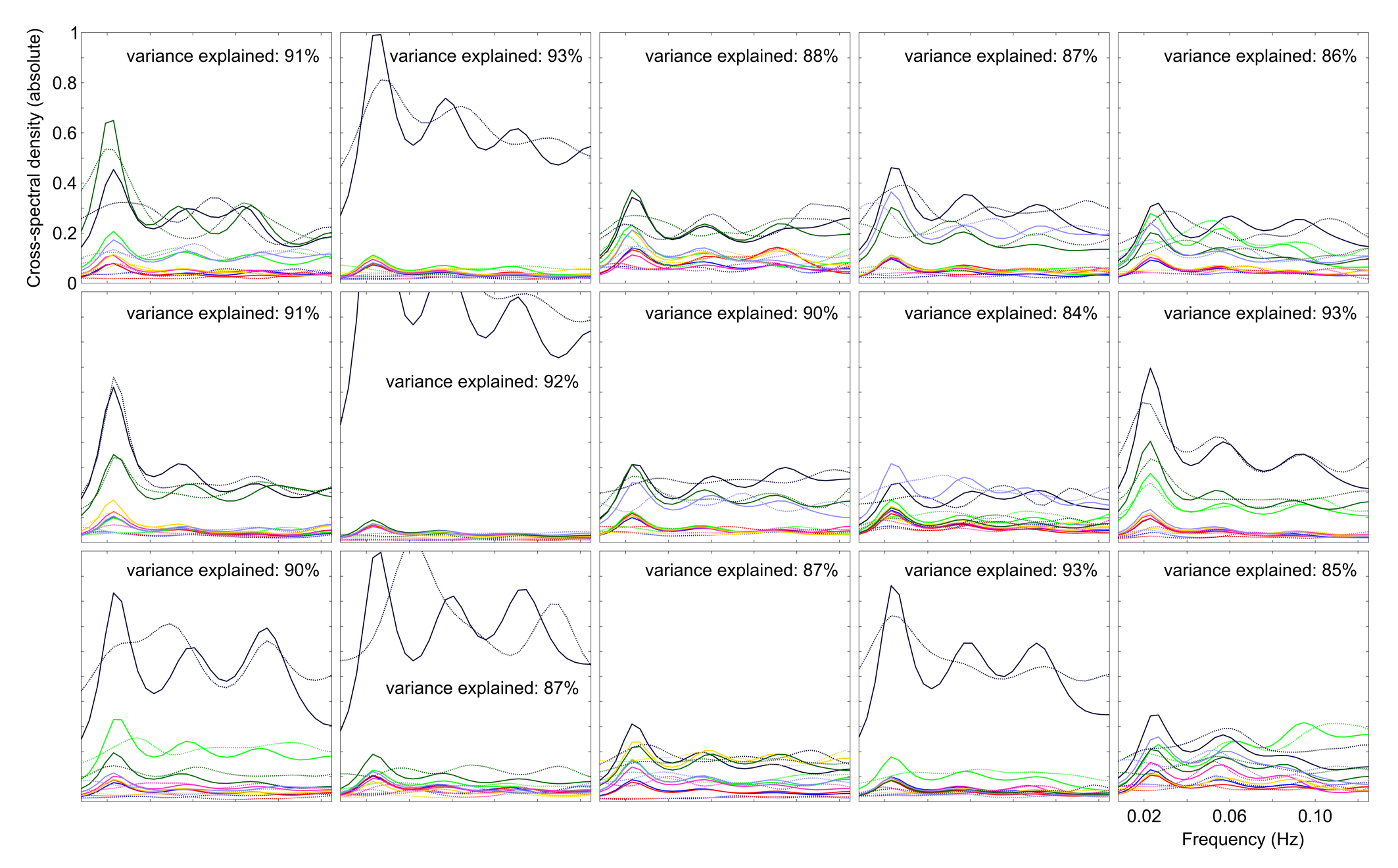

### Supplementary figure 2

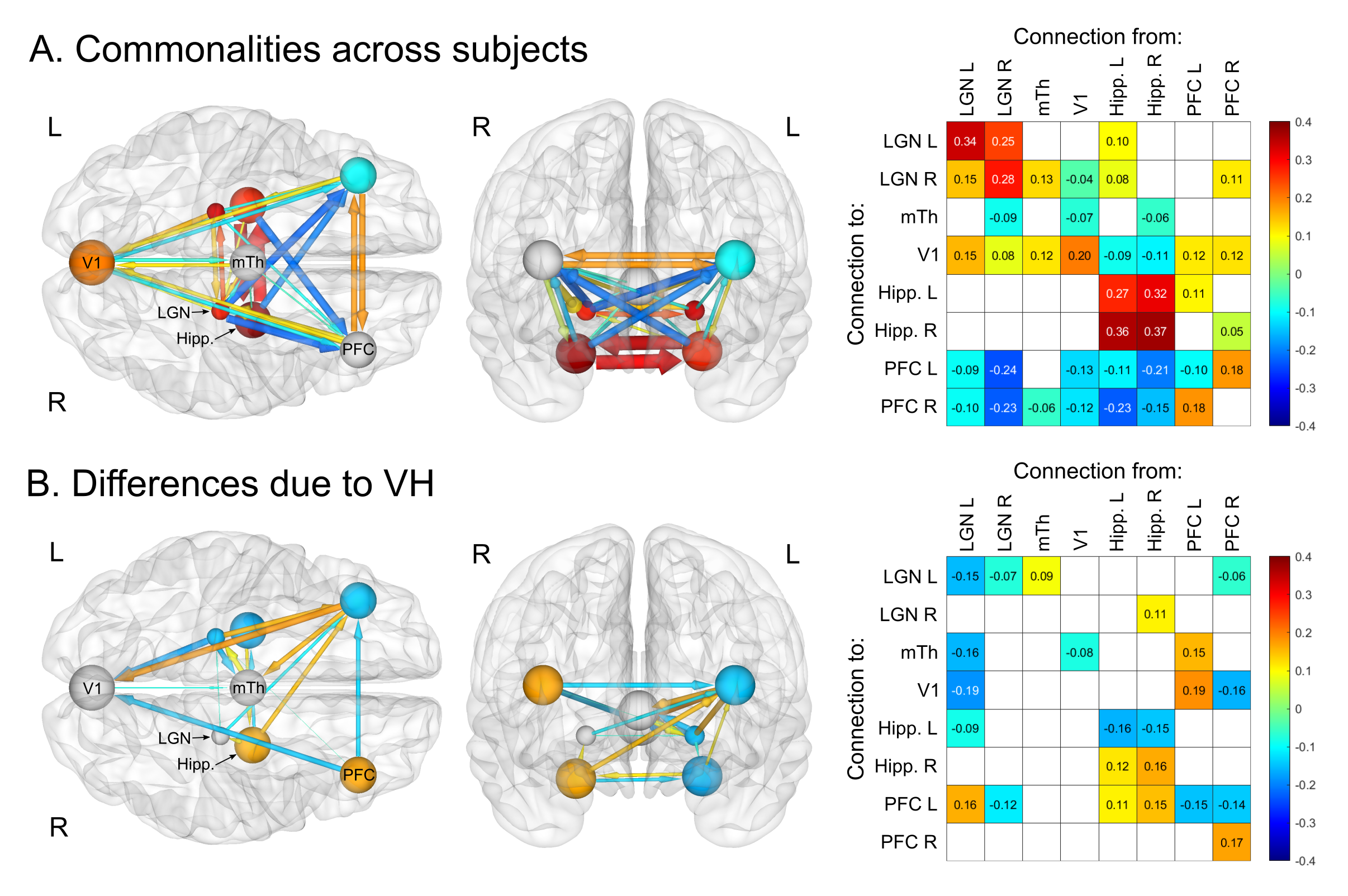
